# Control of cortical cytoskeleton-membrane interaction by RhoA regulates peripheral nerve myelination

**DOI:** 10.1101/2021.11.09.467948

**Authors:** Ana I. Seixas, Miguel R. G. Morais, Cord Brakebusch, João B. Relvas

## Abstract

Bidirectional transmission of mechanical and biochemical signals is integral to cell-environment communication and underlies the function of Schwann cells, the myelinating glia of the peripheral nervous system. As major integrators of “outside-in” signaling, Rho GTPases link actin cytoskeleton dynamics with cellular architecture to regulate adhesion and cell deformation. Using Schwann cell-specific gene inactivation, we discovered that RhoA promotes the initiation of myelination, axonal wrapping and axial spreading of Schwann cells, and is later required to restrict myelin growth in peripheral nerves. These effects are mediated by modulation of actomyosin contractility, actin dynamics and cortical actin-membrane attachment, which collectively couple tensional forces to intracellular signaling that regulate axon-Schwann cell interaction and myelin synthesis. This work establishes RhoA as an intrinsic regulator of a biomechanical response that controls the switch of Schwann cells towards the myelinating and the homeostatic states.

## Introduction

During postnatal peripheral nervous system (PNS) development, Schwann cells (SCs) interact with axons to form the myelin sheath, a remarkable example of membrane-mediated specialized cell-cell communication. This lipid-rich membrane insulates axons to speed nerve conduction and promote long-term axonal integrity. SC development and myelination are coupled multistep, morphogenetic processes. After migrating along axons, perinatally SCs segregate single axons from embryonic bundles, a step called radial sorting (1). The SC in this promyelinating SC-axon unit will then terminally differentiate and radially extend a cytoplasmic wrapping membrane to form the myelin sheath. These events are spatiotemporally regulated, with SCs responding to paracrine and juxtacrine signaling from extracellular matrix (ECM) components and the axonal surface (2).

Rho GTPases are main cytoplasmic integrators of extrinsic mechanical and chemical signals, working as molecular switches that cycle between inactive GDP- and active GTP-bound states. When active, they target numerous effectors that act on the actin cytoskeleton to regulate cellular architecture and diverse cellular functions. Rac1 and Cdc42 are essential for radial sorting (3, 4), and disruption of Rho GTPase signaling is a feature of mouse models with prominent sorting defects (1), including for integrin-linked kinase (ILK) (5), beta1 integrin (4), and for the actin-binding protein profilin-1 (PFN1) (6).

RhoA and its principal kinase effectors ROCK 1 and 2 are master regulators of cortical cytoskeleton organization and contractility, stress fiber formation, and focal adhesion assembly in most cell types (7–9). The Rho subfamily includes the highly homologous RhoA, RhoB and RhoC, all expressed in SCs during myelination (10). Evidence from Ilk, Pfn1 and adhesion G protein-coupled receptor Gpr56 mutants suggests that Rho activity must be tightly controlled for radial sorting to occur (5, 6, 11). Inhibition of ROCK affects SC morphology and results in shorter myelin segments in co-cultures (12). *Rhoa* silencing in rat sciatic nerves causes hypomyelination and affects the motility, proliferation (13) and cAMP-induced differentiation (14) of rat SCs in vitro.

The specific functions of RhoA in myelination remain unknown. Here, we report that SC-specific ablation of *Rhoa* in the mouse delays radial sorting and myelination and, paradoxically, leads to radial hypermyelination later in development. We propose that this dual function of RhoA is mediated by regulation of the structure and contractility of the actin cortical network and its attachment to the plasma membrane, important for bidirectional signaling that regulates axon-glia interaction, wrapping and myelin biogenesis. Together, our results suggest that RhoA modulates a biomechanical response that controls myelin sheath formation, growth and homeostasis.

## Results

### Loss of RhoA in Schwann cells delays radial sorting and the onset of myelination

To study its role in PNS development, *Rhoa* was ablated in myelinating cell precursors from embryonic day (E) 11.5 by expressing Cre recombinase under the control of the Cnp regulatory sequences (15). Recombination of the conditional *Rhoa* allele led to a 90% decrease of RhoA in nerve lysates of postnatal day (P) 5 RhoA mutant mice (*Cnp*-CRE:*Rhoa*^fl/fl^, referred to as RhoA cKO) vs. controls (*Rhoa*^fl/fl^, referred to as CTR) (Figure 1A). RhoA is expressed in SCs (10) and its activity is differentially regulated in the early postnatal sciatic nerve (4). This time window (P0-P5) captures radial sorting, the establishment of 1-1 axon-promyelinating SC units, and the switch to the SC myelinating state and onset of myelination. Ultrastructural analysis of nerves at P1 and P5 showed a decrease in the number of sorted axons in RhoA cKO vs. CTR (Figures 1B, 1C and 1D). This difference diminishes with time (from 22% at P1 to 12% at P5), suggesting that radial sorting is delayed in the RhoA cKO. In line with a sorting impairment, the percentage of axons larger than 1 μm, the threshold axon caliber for myelination, still remaining within bundles at P5 is increased in RhoA cKOs (Figures 1I and 1J). Notably, the number of myelinated axons was decreased in RhoA cKO nerves at P1 (Figure 1E) and P5 (Figure 1F), while the number of promyelinating 1-1 units was similar at P1 (Figure 1G) and increased in RhoA cKO nerves at P5 (Figure 1H), indicating that the rate of the onset of myelination is slower in RhoA mutants.

**Fig. 1.**
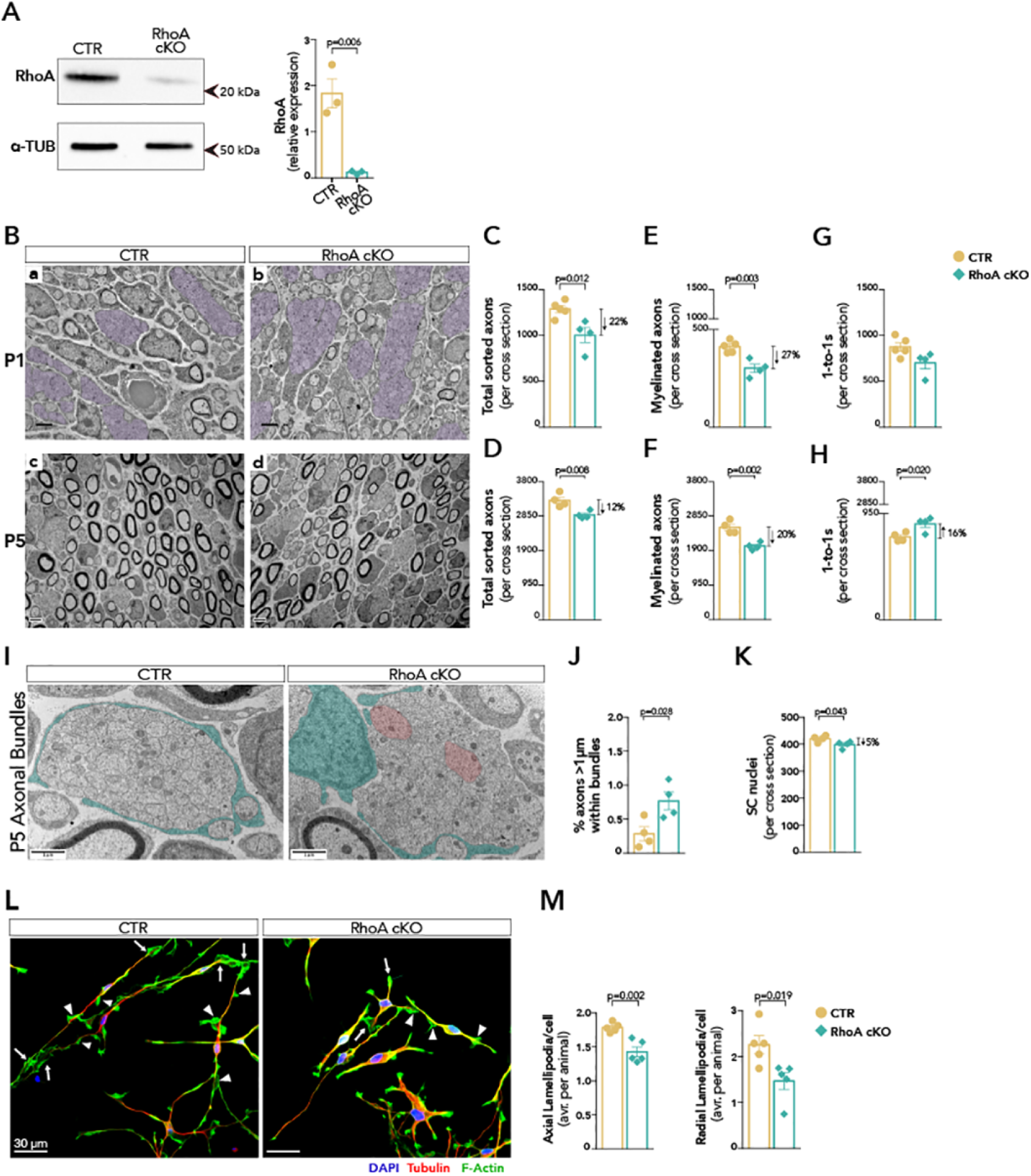
Schwann cell-specific loss of Rhoa delays axonal sorting and the onset of myelination in peripheral nerves. (**A**) Immunoblotting of RhoA in P5 nerve lysates and quantification of signal density relative to α-TUB. n=3 mice/group. (**B**) TEM micrographs of sciatic nerve cross sections from CTR (a,c) and RhoA cKO (b,d) mice at P1 (top) and P5 (bottom). Bundles of unsorted axons are shaded in purple. Scale bar is 2 μm. (**C-H**) Morphometric analysis of whole nerve cross sections at P1 and P5 showing the total number of axons sorted from bundles (C, D) number of myelinated axons (E, F), and axons in 1-1 relationships with SCs (G, H). n=5 or 4 mice/group (C, E, G); n=4 mice/group (D, F, H). (**I**) TEM micrographs showing a bundle of unsorted axons surrounded by cytoplasmic processes of SCs (in green) at P5 and axons larger than 1μm (in red). (**J**) Percentage of unsorted axons larger than the threshold caliber for myelination. Approximately 2000 unsorted axons measured per nerve; and (**K**) Number of SC nuclei in P5 nerve cross sections. n=4 mice/group. (**L**) IF of primary SCs labeling for microtubules (tubulin, red), F-actin (phalloidin, green) and nuclei (DAPI, blue). F-actin labels axial lamellipodia found at the tip of processes (arrow) and radial lamellipodia (arrowhead). (**M**) Quantification of axial and radial lamellipodia in SCs spreading on laminin-2 for 24h post-isolation. At least 50 cells/well were analyzed. n=5 mice/group. All data represented as mean ± SEM and analyzed with unpaired t-test.

### RhoA regulates the timely generation of new SCs and their morphology

We next sought to identify what aspects of SC development underlay the early postnatal phenotypes in the RhoA cKO. Analysis of axonal neuregulin 1(NRG1) type III signaling - a major regulator of PNS myelination with pleiotropic effects on SC development (16) - showed that phosphorylation of ERBB2, a NRG1 receptor in SCs, at P5 was comparable in RhoA cKO and CTR nerve lysates (Figure S1A), suggesting that *Rhoa* ablation does not disrupt NRG1-ERBB signaling in SCs.

Quantification of SC numbers in nerves showed a 5% decrease in the RhoA cKO (Figure 1K), indicating that RhoA is required for the timely generation of new SCs in developing peripheral nerves, a pre-requisite for radial sorting. However, we found no significant changes in SC proliferation (Figure S1B) or survival (Figure S1C) in P5 nerves of RhoA cKOs. RhoA is a major regulator of cytokinesis (17) and loss of *Rho1* in *Drosophila* glia leads to severely decreased glia numbers due to cytokinesis failure (18). So, we hypothesize that, in the RhoA cKO, a subtle delay in cytokinesis may cause SCs to enter differentiation and miss the window for division.

Adequate population of peripheral nerves by SCs requires migration of immature SCs, and disruption of Rho-ROCK signaling impairs the intrinsic motility of SCs in vitro (13, 19). SC migration was analyzed on isolated E13.5 cervical dorsal root ganglia (DRG). At DIV5, and although SCs from DRGs of RhoA cKOs tended to populate more distant segments of the neurite projections, we found no significant difference in migration between groups (Figure S1D) and this is likely not a major contributor to the radial sorting phenotype.

SCs form two types of specialized actin-based protrusions that serve as multifunctional lamellipodia (axial and radial) that establish and stabilize SC-axon contacts and drive wrapping. Defects in SC morphology are linked to radial sorting and/or myelination impairment in several mutants (3–5, 11, 20). SCs from sciatic nerves of RhoA cKO grown in vitro on laminin-2 showed a decrease in both axial and radial lamellipodia compared to SCs of CTR (Figures 1L and 1M), in line with the described effect of ROCK inhibitor fasudil on lamellipodia in cultured mouse SCs (21), suggesting that RhoA-ROCK non-redundantly regulates SC morphology.

Since transcriptional differentiation of SCs is essential for myelination, we analyzed the expression of OCT6/POU3F1 and KROX20/EGR2, master regulators of the SC promyelinating state, and found comparable levels in P5 nerves of CTR and RhoA cKOs (Figures S1E, S1F and S1G). This indicates that in vivo RhoA does not regulate SC differentiation, and is aligned with the reported effect of ROCK inhibition on SC morphology but not on SC proliferation and differentiation (12).

Globally, we observed that RhoA is required to precisely match numbers of SC to axons during radial sorting, and to promote the switch to the myelinating SC state independently of transcriptional differentiation, likely by controlling the formation/stabilization of lamellipodia that mediate axon-SC adhesion.

### Proteomics profiling of early postnatal RhoA cKO peripheral nerves

To identify cellular processes disrupted in the RhoA cKO, we performed quantitative proteomics of P5 sciatic nerves and captured the molecular dynamics of early axon-SC interaction and onset of myelination. Five biological replicates/genotype were analyzed by label-free LC-MS/MS (Figure 2A) and identified 5895 proteins (Table S1). RhoA cKO/CTR abundance ratios were then ranked according to selected post-processing criteria (Figure 2A), yielding 184 proteins that were differentially abundant (DA) (Figure 2B and Table S2).

**Fig. 2.**
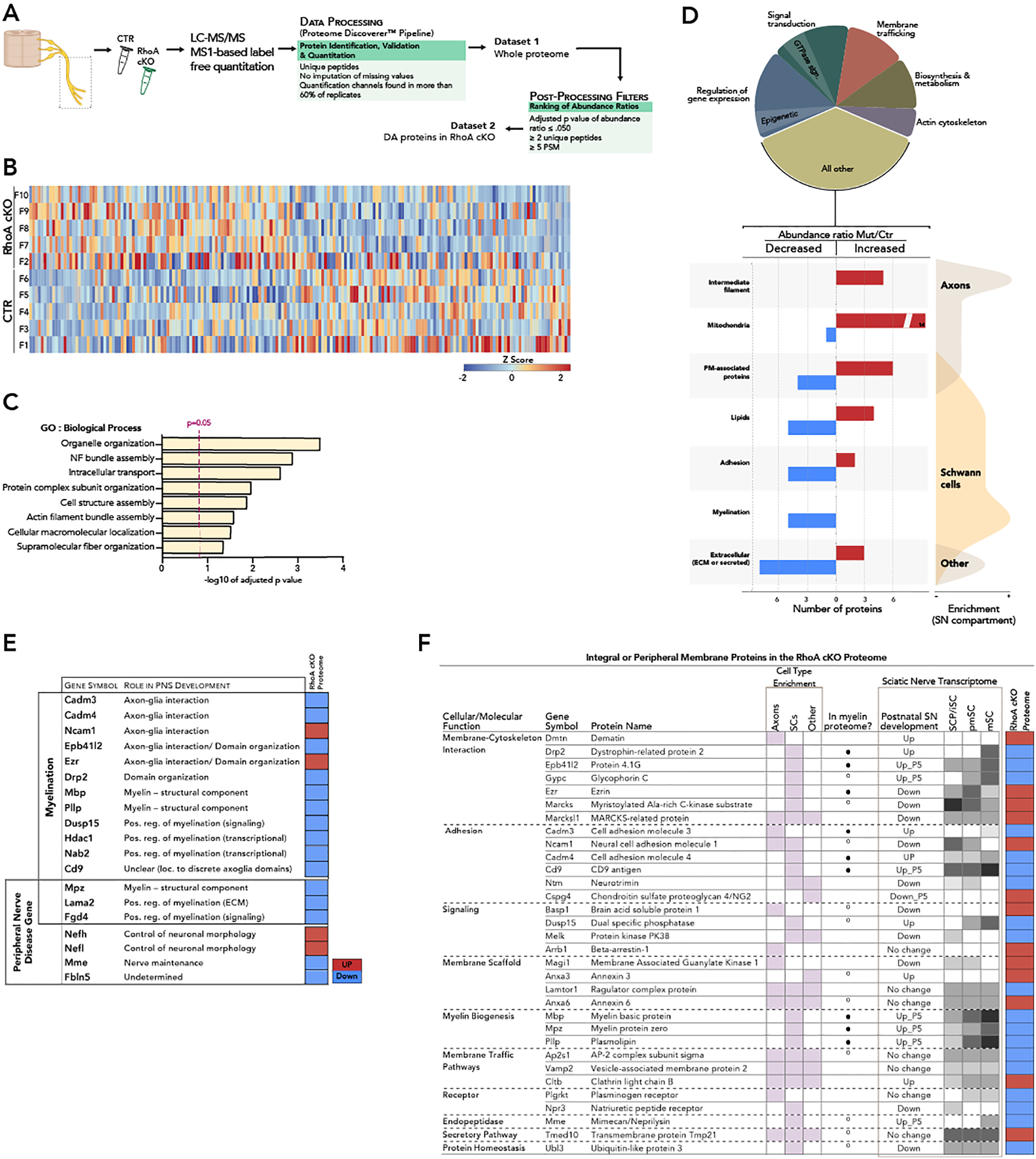
The proteome of early postnatal RhoA cKO peripheral nerves. (**A**) Biological and data processing workflow for the unbiased quantitative proteomics screening of CTR and RhoA cKO sciatic nerves at P5. (**B**) Data matrix of normalized abundances of DA proteins across 5 biological replicates per genotype, represented as a heatmap. Each biological replicate is a pool of 8 nerves from littermates. (**C**) Significantly enriched biological process GO terms of the RhoA cKO sciatic nerve proteome. (**D**) Schematic representation of the functional clustering of 170 DA proteins in RhoA cKO nerves (see also Table S2). The pie chart displays all functional clusters with more than five DA proteins, and below a breakdown of predicted compartment-specific contributions to selected functional clusters. (**E**) List of RhoA cKO DA proteins involved in myelination and/or peripheral nerve development/maintenance. (**F)** Membrane proteins in the RhoA cKO proteome. Most are expressed by SCs and sciatic nerve transcriptome data predicts a differential expression during postnatal development. Circle symbol - found in two (black) or one (white) myelin proteome datasets. Shades of grey refer to relative abundance/timepoint (lighter tones refer to lower abundance, darker tones to higher abundance). DA - differentially abundant; PM - plasma membrane; SN - sciatic nerve; SCP - SC precursors; iSC - immature SCs; mSC - mature SCs

Next, we performed a gene ontology (GO) analysis (Figure 2C) and further manually annotated the DA protein dataset into 12 functional clusters of 170 proteins (Figure 2D). The sciatic nerve is highly enriched in SCs but also has fibroblasts and axons, so we searched scRNAseq databases (10, 22, 23) to predict cell type-predominant protein expression. Seven clusters tended to be globally inhibited (myelination, adhesion and ECM) or upregulated (e.g. intermediate filaments and mitochondria of axons) in specific cell compartments of RhoA cKO peripheral nerves, denoting that loss of *Rhoa* in SCs impacts global nerve physiology at the onset of myelination (Figure 2D). DA proteins known to positively regulate PNS myelination were predominantly decreased, including myelin components, axon-glia interface signaling and adhesion molecules, and proteins encoded by peripheral disease genes (Figure 2E).

Remarkably, nearly 20% of the RhoA cKO proteome are integral or peripheral membrane proteins, including cytoskeleton-membrane complex proteins important in cellular processes dysregulated in the RhoA cKO, such as signaling via receptors, adhesion and membrane trafficking (Figure 2F). Most (24/32) of these proteins are found in SCs, predicted to be differentially expressed during postnatal sciatic nerve development, and 67% (20/32) have been previously identified in the peripheral myelin proteome (24, 25). A subset of these membrane-associated proteins are well-characterized axo-glia interface cytoskeletal and adhesion molecules, and modulators of actin cytoskeleton-membrane interaction.

Altogether, these observations point towards altered molecular properties of the SC surface in the RhoA cKO, which likely impede on bidirectional signaling that regulates axon-SC and ECM-SC interaction at the onset of coordinated axonal wrapping and myelin biogenesis.

### Dysregulation of cytoskeletal signaling and cortical tension in SCs from the RhoA cKO

The proteomics findings predict that RhoA modulates cytoskeletal organization and its interaction with the peripheral membrane of SCs, important for the transmission of cellular force. So, we asked whether mechanical properties of the SC cortex in the RhoA cKO were changed. To measure cortical cytoskeletal stress in live SCs, we expressed a genetically-encoded force sensitive α-actinin FRET probe (actinin-sstFRET) (27) with a donor/acceptor fluorophore pair separated by a spectrin linker inserted into actinin, such that increased tension pulls the fluorophores apart and yields lower FRET ratios. Transfected, steady-state SCs did not show obvious morphological perturbations and displayed increasing tension towards the cell’s lamellipodia (Figure 3A). We observed increased whole-cell FRET ratio in SCs from RhoA cKO nerves vs. CTR (Figure 3A and 3B), indicating lower tension, suggesting that in SCs, RhoA is a major regulator of internal stresses in the cortical cytoskeleton, which are simultaneously responsive and determinant of physical properties that generate force and drive cell deformation.

**Fig. 3.**
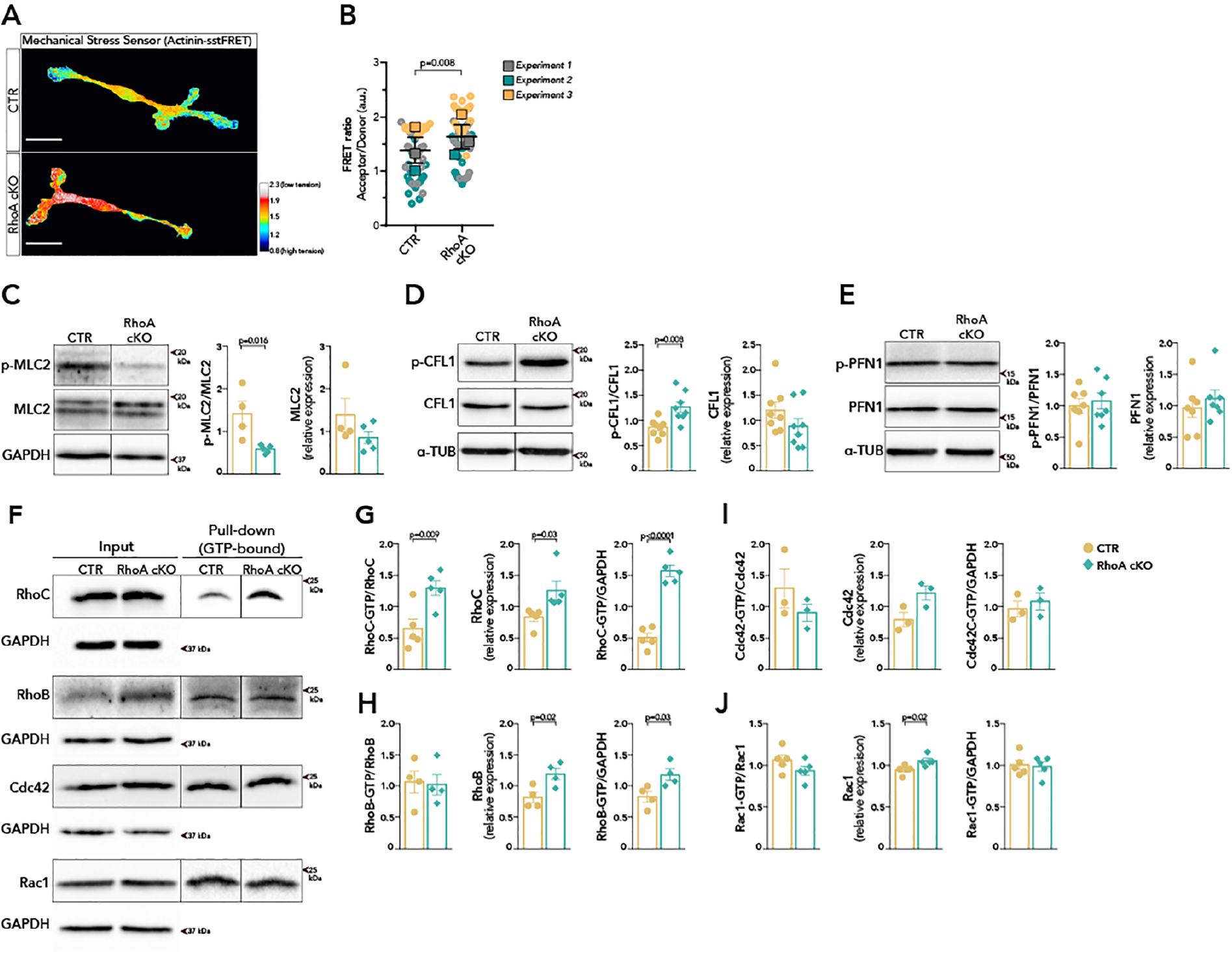
Disrupted cortical tension, actomyosin contractility and actin dynamics in SCs from RhoA cKO mice. (**A**) Pseudocolored images displaying pixel value of FRET ratio in primary mouse SCs expressing the actinin-sstFRET tension sensor. (**B**) Quantification of FRET ratios. The SuperPlot (26) shows superimposed data from 60 cells/genotype with mean FRET ratio ± SEM of n=3 independent experiments. Scale bar is 25 μm. (**C-E)** Immunoblotting of phosphorylated and total MLC2 (C), CFL1 (D) and PFN1 (E) in P5 nerve lysates. Quantification of signal density relative to total protein, α-TUB or GAPDH. n=4 or 5 (C), n= 8 (D), n=7 (E) mice/group. (**F**) Analysis of GTP-bound (active) levels of Rho GTPase family proteins by pull-down assay of P5 nerve lysates. (**G-J**) The activated fraction was calculated by normalizing signal densities of GTP-bound to total Rho GTPase. Total levels of Rho or Rho-GTP is relative to GAPDH. n=5 (G), n=4 (H, J), n=3 (I) biological replicates. Data represented as mean ± SEM and analyzed with: unpaired t-test (D, E, G-J), Mann-Whitney (C) or paired t-test (B).

We next sought to identify downstream Rho-ROCK effectors that are modulating the cortical actin cytoskeleton in SCs. The actomyosin complex is the central regulator of contractile forces and its myosin light chain 2 (MLC2) subunit is a direct target of ROCK (28), including in SCs (12). Levels of phosphorylated (p-) MLC2 were decreased in P5 nerve lysates of RhoA cKO vs. CTR (Figure 3C), indicating diminished cortical actomyosin contractility. Analysis of the actin-severing and depolymerizing protein Cofilin-1 (CFL1), which can be inactivated by Rho-ROCK-LIM kinase, showed that levels of p-CFL1 were increased in P5 RhoA cKO nerves (Figure 3D), suggesting decreased actin filament turnover. Previous studies in vitro suggest that activity of myosin II and cofilin is required for myelination (29, 30). In line with those proposed models, disruption of actomyosin contractility and actin turnover upon loss of RhoA in SCs are probably major contributors to the impaired axonal wrapping and onset of myelination in the RhoA cKO. Surprisingly, we found no changes in levels of p-PFN1, an actin-binding protein that positively regulates radial sorting and myelination and can be modified by ROCK (6), in P5 nerves of RhoA cKO vs. CTR (Figure 3E). This observation, together with the increased levels of p-CFL1, suggests an uneven disruption of the Rho-ROCK pathway, or a compensatory activation of other kinases, when Rhoa is ablated in SCs. Rho GTPases can cross-regulate each other (31) and have promiscuous interactions with many effectors (32). Using a Rho-GTP pull down assay of P5 nerve lysates, we determined that the expression and total active levels of both active (GTP-bound) RhoB and RhoC were upregulated in RhoA cKOs (Figure 3F, 3G, 3H). We next used a similar approach to analyze active fractions of Rac1 and Cdc42, since both are important for PNS myelination and Rac1 is a main regulator of radial lamellipodia formation in SCs (3, 4). We did not detect significant changes in levels of active Rac1 and Cdc42 in RhoA cKO nerves (Figure 3F, 3I, 3J).

### RhoA regulates myelin sheath growth and homeostasis

We next assessed if the myelination program resumes normally in the RhoA cKO PNS later in development. Ultra-structural and g-ratio - a normalization of axon to myelinated fiber diameter - analyses of sciatic nerves at P15 and P25 showed thinner myelin sheaths in RhoA cKO vs. CTRs (Figures 4A and S2B). Similar findings were observed in SC-specific *Rhoa* mutants generated using a different Cre-expressing line (Figures S2A and S2C), the Dhh-Cre mouse (33). This radial hypomyelination can result from the delayed onset of myelination, or reflect a continuous slower rate of myelin growth.

**Fig. 4.**
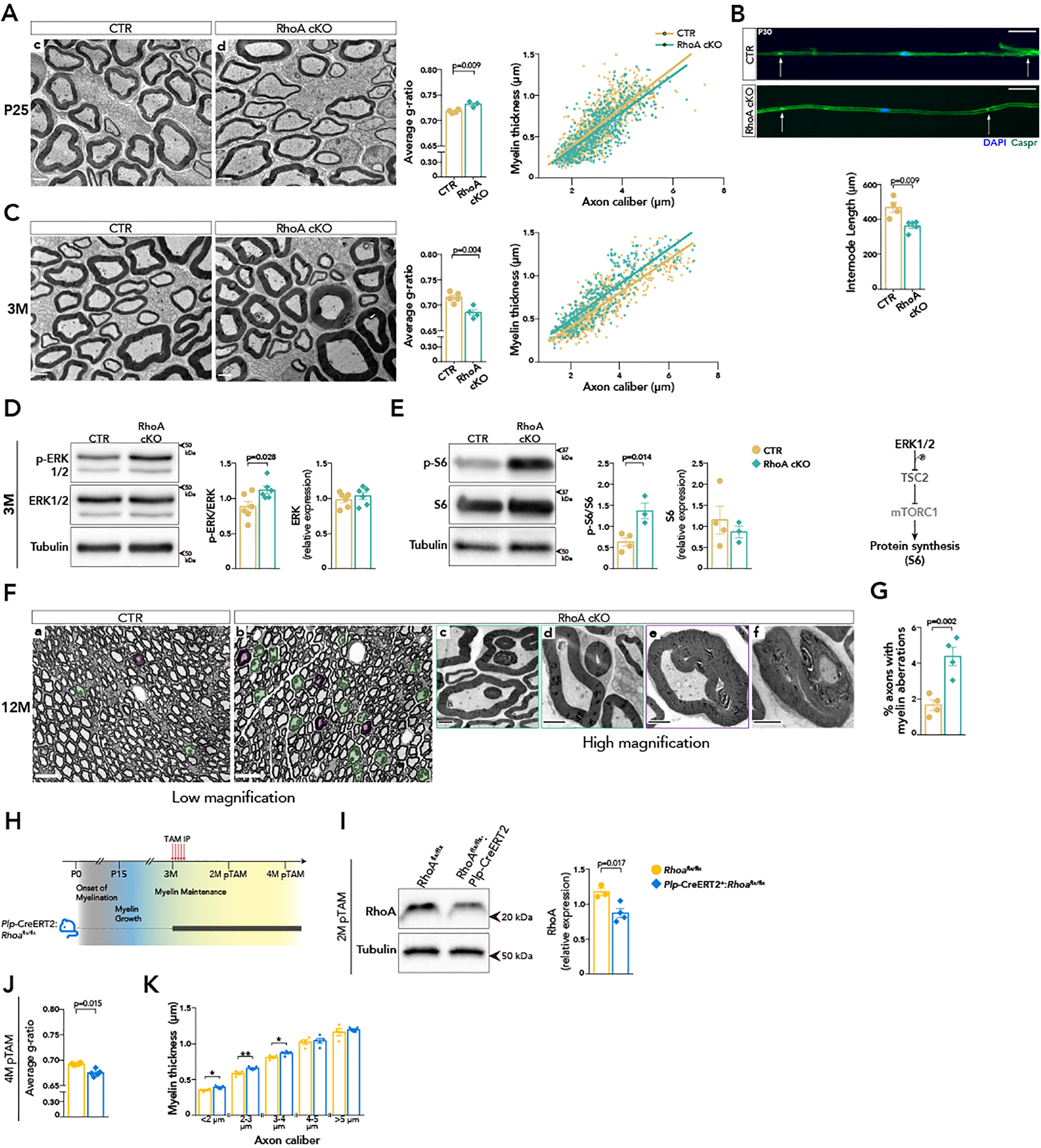
RhoA regulates myelin sheath growth and homeostasis in peripheral nerves. (**A, C**) TEM micrographs of sciatic nerve cross sections from CTR and RhoA cKO mice at P25 (A) and 3M (C). Bar graphs show average g-ratio and scatter plots show distribution of myelin sheath thickness as a function of axon caliber. At least 100 myelinated axons measured/animal. n=4/3 (A) and n= 5/4 (C) mice/group. Scale bar is 2 μm. (**B**) IF of teased fibers labeled for the paranodal protein CASPR (green, arrows) and nuclei (in blue) and quantification of internode length. n=4/5 mice/group. Scale bar is 50 μm. (**D, E**) Immunoblotting of phosphorylated and total ERK1/2 (D) and S6 (E) in 3M nerve lysates. Quantification of signal density relative to total protein or α-TUB. n=6 (D), n=4/3 (E) mice/group. (**F**) Images of semithin cross sections from CTR (a) and RhoA cKO (b) sciatic nerves at 12 months, showing abnormal myelin profiles (pseudocolored). TEM micrographs of RhoA cKO nerves show infolded (c) and outfolded (d) myelin loops, tomacula (e), and large outfolds from unstructured myelin (f). Scale bar is 20 μm (low mag) and 2 μm (TEM). (**G)** Quantification of abnormal myelin profiles. n=4 mice/group. (**H**) Conditional deletion of Rhoa in mature SCs at 3M. (**I**) Immunoblotting of RhoA in sciatic nerves 2 months pTAM and quantification of signal density relative to α-TUB. n=3/4 mice/group. (**J**) Average g-ratio and (K) Categorization of myelin sheath thickness as a function of axon caliber in sciatic nerves 4M pTAM. (J, K) n=4/5 mice/group. *p<0.05, **p<0.01. At least 100 myelinated axons measured/animal. All data represented as mean ± SEM and analyzed with unpaired t-test.

It has been shown that chemical inhibition of ROCK leads to shortened myelin segments in co-cultures (12, 34), suggesting that Rho-ROCK may participate in internode formation. Single fibers teased out from P30 nerves immunolabeled for the paranodal axonal protein CASPR1 showed that internodes were shorter in fibers of RhoA cKO vs. CTR mice (Figure 4B), indicating that RhoA also regulates the axial growth of peripheral myelin sheaths. After developmental myelin biogenesis, myelinating SCs switch to a homeostatic state to maintain sheath stability. Morphometric analysis of nerves at 3 months (3M) showed increased myelin thickness (hyper-myelination) in RhoA cKOs across all axon calibers (Figure 4C).

Restricting myelin growth at the end of myelination is an active process that in the PNS is modulated, at least in part, by the PI3K-AKT (35) and the MAPK MEK-ERK1/2 pathways (36). Downstream, they target the metabolic mTOR pathway (37), which drives myelin synthesis. Mutants with hyperactivated AKT (35), mTOR complex 1 (mTORC1) (38) or ERK1/2 (36, 39) show radial hypermyelination. While we detected no changes in AKT activation, or levels of its inhibitor PTEN, (Figures S4A and S4B), levels of p-ERK1/2 and p-S6 ribosomal protein, a substrate of mTORC1-S6K that upregulates protein synthesis, were increased in the hypermyelinated nerves of 3M RhoA cKOs (Figures 4D and 4E). Hence, RhoA is required to limit developmental myelin growth, by repressing an ERK-mTORC1 signaling cascade that controls myelin synthesis.

Proportionality between myelin thickness and axon caliber is a feature of normal peripheral nerve physiology and the presence of excessive myelin is seen in human neuropathies. Upon analysis of 12M nerves, we found increased aberrant myelin profiles in the RhoA cKO, including myelin infolds and outfolds, tomacula (focal myelin swellings) and unspecific myelin degeneration (Figures 4F and 4G), indicating that RhoA is required to maintain the homeostasis of peripheral myelin sheaths.

A previous study described that levels of active RhoA-GTP increase in adult sciatic nerves (4). So, we questioned if RhoA participates in myelin maintenance. To uncouple from its roles in development, we recombined *Rhoa* in adult SCs using the inducible *Plp*-CreERT2 mouse strain (40) and tamoxifen administration at 3 months, after the critical period of developmental myelination (Figure 4H). Two 2 months later (2M pTAM) we confirmed loss of RhoA in *Plp*- CreERT2:*Rhoa*^fl/fl^ nerve lysates (Figure 4I). At 4M pTAM, an ultrastructural analysis showed increased myelin thickness predominantly in small and medium caliber axons of mutant nerves (Figures 4J and 4K), indicating that *Rhoa* ablation reactivates the growth of adult myelin sheaths. Altogether, these data show that RhoA is required to establish and maintain the homeostatic state of the myelinating SC, and limit myelin growth in adult peripheral nerves.

## Discussion

In this study, we provide evidence that the small GTPase RhoA is required to initiate myelination and drive axonal wrapping, and to curb myelin growth in peripheral nerves. RhoA modulates the organization and contractility of the cortical actin network and its attachment to the plasma membrane, thus coupling mechanical and biochemical signaling that controls the SC switch to the myelinating and, later, homeostatic state.

It is well established that Rho-ROCK signaling downstream of integrin and G protein-coupled receptors modulates radial sorting, and its activity must be finely tuned as both hyperactivated or decreased Rho is detrimental for this step of myelination (5, 6, 11). We have now directly addressed this point and determined that RhoA is required for timely radial sorting. We propose that RhoA participates in the generation of new SCs necessary to match the number of axons to be sorted, and in addition modulates lamellipodia formation/stabilization in SCs, important for adhesion to both axons and the ECM. An important consideration is that the sorting delay in the RhoA cKO is accompanied by an increase in active RhoB and RhoC. RhoC shares high homology with RhoA and can target common effectors also via ROCKs (32, 41). This raises the hypothesis that amplified Rho-ROCK, especially via RhoC, can be offsetting or disrupting the targeting of some effectors in RhoA cKO SCs. While it is evident that loss of RhoA is not compensated, this observation leaves interesting questions as to whether RhoA and RhoC may regulate specific effectors in SCs during radial sorting by interacting differentially with ROCK1/2 at specific subcellular locations, similar to what has been described for cofilin inactivation in actin-rich structures of cancer cells (42).

A prominent phenotype of the RhoA cKO is a delay in myelination independent of SC transcriptional differentiation. This favors a hypothesis that a biomechanical process, uncoupled from differentiation, drives wrapping and myelin biogenesis, as it has been advanced by colleagues in the field (20, 43–45). Targeted membrane extension, deformation, and adhesion require generation of intracellular forces by way of actin filament dynamics and modulation of cortical cytoskeleton organization, which is enriched in myosins, crosslinkers and membrane-anchoring proteins (46, 47). We now provide a clearer picture of the molecular mechanisms that balance intracellular forces that drive myelin growth. In SCs, RhoA induces cortical actomyosin contractility and its crosstalk with actin filament turnover by modifying cofilin activity and, importantly, targets the attachment of the cortical cytoskeleton to the membrane. Of note, the proteomics analysis identified Ezrin (EZR), a regulator of membrane-actin cortex adhesion and component of paranodal SC membranes (48), and Gamma-Tropomyosin (TPM3), an actin-binding contractile protein highly expressed in SCs (10), both increased in the RhoA cKO. Interestingly, TPM3 regulates the mechanical access and activity of other actin-binding proteins, such as myosin II and cofilin, (**?** ), to actin filaments to determine cell cortex stiffness and adhesion (49). It is thus probable that RhoA participates in the generation but also the propagation of cellular forces and contractility, which is highly dependent on the structure of the actin filament network. We propose that these mechanical properties determine the molecular makeup of the SC surface, impacting bidirectional signaling that modulates axon-glia interaction, myelin production and basal lamina composition, all downregulated in the RhoA cKO proteome, and the transition to the myelinating SC state.

RhoA has a dual function in myelination, since adult nerves of RhoA cKO show radial overgrowth of myelin associated with hyperactivation of a MAPK ERK-mTORC1 signaling axis. Maintenance of the SC homeostatic state and control of myelin thickness appears to be a bona fide function of RhoA, uncoupled from earlier developmental defects, since ablating Rhoa specifically in mature SCs reactivates myelin growth. Similar dual functions have been described in models of mTORC1 and PI3K-AKT hyperactivation, although in these models the myelination impairment is linked to deficient SC transcriptional differentiation (38), which does not occur in the RhoA cKO. ERK activation by phosphorylation has been described to depend on Rho-ROCK in some cell types (50, 51) and on actomyosin-driven tensile forces at stress fibers, components of focal adhesions that act as transduction platforms for biochemical signaling (52). Thus, we propose that RhoA is an important molecular determinant for the formation and maintenance of myelin by coupling mechanical signaling to specific intracellular pathways at different phases of the myelin sheath lifetime.

## Materials and Methods

### Animals

The mouse strain with a floxed *Rhoa* allele (53), the transgenic line expressing Cre recombinase under the control of the Cnp (2’,3’-cyclic-nucleotide 3’- phosphodiesterase) promoter (15) and the resulting conditional knockout mice, all in a C57BL/6 background, were housed in cages with environmental enrichment in SPF conditions, kept in a 12-h light and dark cycle, and fed Tecklad 2014S chow ad libitum. All animal procedures, 3Rs and ethical considerations were revised and approved by the i3S Animal Welfare Body and the national authority (DGAV license 024108), which apply the EU directives for animal research. To inactivate *Rhoa* specifically in myelinating cells, *Rhoa*fl/fl animals were crossed with *Cnp*-Cre:*Rhoa*^wt/fl^ or *Cnp*-Cre:*Rhoa*^fl/fl^ to generate *Cnp*-Cre:*Rhoa*^fl/fl^ and *Rhoa*^fl/fl^ experimental animals, here designated as RhoA cKO and CTR, respectively. Genotyping for the Rhoa floxed and the Cnp-Cre knock-in alleles was performed by PCR of genomic DNA isolated from ear or tail using previously published primer combinations (15, 53). For experiments with animals younger than P15, we favored the use of littermates to control for differences in postnatal developmental progress among litters. Animals of both sex were used in all experiments. To inactivate Rhoa specifically in adult myelinating cells, the floxed strain was crossed with a *Plp*-CreERT2 line (40). Recombination was induced by intraperitoneal administration of tamoxifen (2 mg) prepared in a 9:1 oil:ethanol mix (20 mg/ml) for 5 consecutive days. To generate the conditional knockout under the control of the *Dhh* (Desert hedgehog) promoter, a similar breeding strategy was followed using the *Dhh*-Cre strain (33). Tissue collection from *Dhh*- Cre:*Rhoa*^fl/fl^ animals was performed with the approval and in strict accordance with the guidelines of the Zurich Cantonal Veterinary Office.

### Ultrastructural and morphometric analyses

For electron microscopy (EM) analysis, deeply anesthetized animals were transcardially perfused with fixative (3% glutaraldehyde and 4% paraformaldehyde (PFA) in 0.1 M phosphate buffer), sciatic nerves dissected and post-fixed for 24-48h at 4°C. To prepare for sectioning, tissue was put in 1% osmium tetroxide (EMS), dehydrated in serial dilutions of increased acetone concentration and embedded in Spurr resin (EMS). Ultra-structural analysis was performed on ultrathin sections (65 nm) imaged in a Jeol JEM 1400 TEM. Measurements of myelin thickness and axon diameter were performed as described in (54) on 100-200 fibers/animal sampled from 5-7 randomly acquired images of the nerve. Quantification of aberrations at 1 year was performed on half-nerve reconstitutions of semithin sections (0.5 μm) stained with toluidine blue and analyzed at low magnification in a Nikon Ti microscope. All image analysis and quantifications were performed in Fiji (55, 56) using the available standard selection and measurements tools.

### Preparation of frozen sections and teased fibers from sciatic nerves

For immunofluorescence analysis of tissue, deeply anesthetized animals were transcardially perfused with 4% PFA/PBS, sciatic nerves dissected, post-fixed for 12-16h at 4°C and then moved to a 20% sucrose solution overnight. To prepare for sectioning, nerves were embedded in OCT (Tissue Tek), frozen and cut into 10-12 μm sections on a Leica CM 3050S cryostat. To prepare teased fibers, sciatic nerves were dissected, fixed in 4% PFA/PBS for 4h at 4°C and then moved to PBS. After removal of the perinerium,1 cm pieces were divided into fascicles and individual fibers were teased apart with a fine needle on APES-coated microscope slides.

### Isolation and transfection of mouse Schwann cells

Sciatic nerves were dissected from P3-P4 animals and stripped of the perineurium in HBSS with penicillin-streptomycin (Gibco). To isolate SCs, we proceeded as described (54) with minor modifications. Briefly, tissue was enzymatically digested in trypsin (1.25 mg/mL) and collagenase type I (2 mg/mL) (both from Sigma) for 45 minutes, followed by gentle mechanical dissociation in cDMEM (DMEM GlutaMAX, 10% FBS, Gibco). Cells were seeded on 5 μg/ml laminin 2-coated glass coverslips in cDMEM medium, which was replaced on the next day by SC medium (DMEM/F-12 [Gibco], N2 supplement [Gibco], 10 ng/ml Heregulin-β1 (EGF Domain) [Sigma] and 2.5 μM Forskolin [Sigma-Aldrich]). For transfection, after the mechanical dissociation step and before seeding, the cell pellet was gently resuspended in 20 μL of P3 nucleofection solution, 1 μg plasmid DNA was added and cells electroporated in a 4D-NucleofectorTM System (Lonza) following the manufacturer’s instructions.

### Immunofluorescence, TUNEL and EdU assay

For the TUNEL (terminal deoxynucleotidyl transferase dUTP nick end labelling) assay, tissue sections were incubated for 1h in blocking buffer (10% normal goat serum [NGS], 0.1% BSA and 1% Triton X-100 [TX] in PBS) and equilibrated for 10min in TdT buffer (30 mM Tris-HCl, 140 mM sodium cacodylate trihydrate 1mM cobalt (II) chloride hexahydrate). Integration of biotin-16-dUTP (Roche) was performed for 1.5h at 37°C with TdT enzyme (Roche) and dATP (Promega) in TdT buffer, and the detection with streptavidin conjugated with AlexaFluor-568. For detection of proliferative cells, the Click-iT EdU (5-ethynyl-2’-deoxyuridine) Alexa Fluor (AF) 568 imaging kit (Molecular Probes) was used in accordance with the manufacturer’s instructions. Teased fiber preparations were permeabilized in 0.25% TX-PBS for 30 min and blocked for 1h in 1% NGS/1% horse serum (Gibco)/1% HEPES/PBS. Incubation with an antibody against contactin-associated protein (CASPR) was performed overnight at 4°C in blocking solution, followed by an incubation with antimouse-AF488. For SCs grown in vitro, fixation was performed in microtubule protecting fixative buffer (65 mM PIPES, 25 mM HEPES, 10mM EGTA, 3 mM MgCl2, 4% PFA) for 15 minutes, followed by blocking for 1h in 10% NGS-PBS, and an overnight incubation at 4°C with an antibody against Tubulin in 10% NGS-PBS. A secondary antibody against mouse and 658-Phalloidin (1:50, Invitrogen) were diluted in blocking solution and added to the cells for 1h. All preparations were stained with 4’,6’-diamidino-2-phenylindole (DAPI, Invitrogen) for 10 minutes to label nuclei and mounted with Fluoroshield medium (Sigma). Imaging was performed on a Leica DMI6000 equipped with an Orca Flash 4.0 v2.0 camera (Hamamatsu) using a 40x/0.60 objective, or a Nikon Ti equipped with an iXon888 camera (Andor) using a 40x/0.95 objective, epifluorescence microscopes. All image analysis and B&C adjustments were performed in Fiji and the “Cell Counter” plugin was used for quantifications.

### SC migration assay in DRG explants

Cervical dorsal root ganglia (DRG) were isolated from E13.5 mouse embryos, when immature SCs migrate along the axonal tracts of spinal nerves (57). Dissection was performed in HBSS with Pen/Strep/Amph (Gibco). Explants were placed on coverslips previously coated with matrigel (Corning) diluted 1:5 in Neurobasal (Gibco) and grown for 5 days in the following medium: Neurobasal, B27, Glutamax and nerve growth factor (10 μg/mL). Embryos were genotyped using tail tissue. Fixation was performed in 4% PFA-PBS for 15 minutes, followed by an incubation in 0.1M glycine, permeabilization in 0.25% TX-0.1% Tween-PBS, and blocking for 1h in 2% BSA-1% normal donkey serum-0.1% Tween-PBS. Coverslips were incubated overnight at 4°C with antibodies against SOX10 and ß3-Tubulin in blocking buffer and stained with DAPI. Imaging was performed on a Leica DMI6000 microscope using a 20x/0.40 objective. Tilescan images of whole DRGs were acquired using LasX Navigator. Semiautomatic counting of SCs at fixed distances from the explant was performed using a modified Sholl analysis macro in Fiji (code available at https://github.com/mafsousa/ALM_BIA-scriptingtools/blob/main/CreateBandsQuadrants.ijm). Briefly, 20 concentric rings spaced by 200 px were overlayed on the image and SOX10/DAPI positive cells were counted. This quantification was performed by quadrant, to exclude overlapping neurites from neighboring DRGs. The final percentage of SCs as a function of distance to explant is the average of quadrants analyzed/DRG.

### Live FRET analysis

Mouse SCs were transfected with the PEG-Actinin-C-sstFRET plasmid, a gift from Fred Sachs (Addgene plasmid 61101; RRID:Addgene_61101) (27), encoding a force sensitive FRET-based probe, and grown on laminin-2 coated 2 well μSlides (Ibidi) for 60h. Two hours before live cell imaging, fresh SC medium prepared in FluoroBrite-DMEM was added to the cells. Imaging was performed on a Leica DMI6000 with temperature and CO2/O2 control, equipped with an Orca Flash 4.0 v2.0 camera (Hamamatsu), a mercury metal halide lamp with an integrated light attenuator, external excitation and emission filter wheels, and using a HCX PL APO CS 63x/1.30 GLYC objective. For dual-emission ratio imaging, we used BP 427/10 and BP 504/12 excitation filters for CFP and YFP, respectively, a GC dichroic cube (440/520), emission filters BP 472/30 for CFP and BP 542/27 for YFP, and exposure time of 0.8s with the CCD camera binning set to 2×2 at a 16-bit depth. Ratiometric FRET was calculated as acceptor/donor, as described in (Meng and Sachs, 2011). Image analysis was performed semi-automatically using an in-house developed macro for Fiji that allows for batch-processing of files (code available at https://github.com/mafsousa/2DFRETratiometrics). The main workflow consists of the following steps: 1) a preprocessing stage including shading correction and background subtraction (either by using a background image or by subtracting user-defined background mean intensity values); 2) cell segmentation using a user-selected channel, the best threshold algorithm and an option for user-dependent refinement; and 3) ratiometric analysis by dividing selected preprocessed channels in the segmented cell with final ratio images represented with a Royal LUT.

### Preparation of total tissue lysates and immunoblotting

Dissected sciatic nerves were stripped of perineurium and epineurium, crushed on dry ice and homogenized with a chilled pestle in lysis buffer (50 mM Tris pH 8.0, 150 mM NaCl, 1% NP-40, 1 mM EDTA, 0,5% sodium deoxycholate, 0,1% SDS, 1% protease and phosphatase inhibitor cocktails [Sigma]). Samples were left on ice for 30 min, centrifuged at 13000 xg to clear the lysate and stored at −80°C. For immunoblotting, protein extracts were resolved by SDS-PAGE in 10% or 12% polyacrylamide gels, transferred onto a nitrocellulose membrane that were then blocked for 1 h at room temperature in 5% (w/v) milk or BSA in TBS. For analysis of ERBB2, protein extracts were prepared in lysis buffer with 2% SDS and resolved in a 4-12% gel (Biorad). Membranes were probed overnight at 4°C, and for 1 h at room temperature for primary and secondary antibodies, respectively, and signal detection was performed with SuperSignal West Pico Chemiluminescent Substrate (ThermoFisher) on a ChemiDoc XRS digital system (Biorad). Analysis of signal intensity was performed in Fiji using the “Analyze-Gels” menu and from images obtained within the linear range of detection. Relative expression levels are calculated from normalization to α-tubulin or GAPDH.

### Rho-GTP pull down assay

Sciatic nerves (pooled from 3 littermates/biological replicate) lysates were prepared as described above in a modified immunoprecipitation buffer (10% glycerol, 50 mM Tris-HCl pH 7.4, 100 mM NaCl, 1% NP-40, 2 mM MgCl2 and protease inhibitor cocktail [Sigma]). 10% of lysate volume was reserved to determine total protein amounts, and the remaining protein mixture was immunoprecipitated with the respective bait substrate (GST-tagged p21-activated kinase-binding domain [GST-PAK-PBD], provided by J. Collard, NKI, The Netherlands, for Rac1 and Cdc42; and GST-tagged rhotekin Rho-binding domain [GST-RBD], purchased from Millipore, for Rho proteins) that was immobilized on Glutathione Sepharose beads (GE Healthcare). Bait-couple beads were incubated with lysates incubated with overnight at 4°C, washed with 10 packed volumes of lysis buffer, and bound proteins were eluted in Laemmli buffer. Immunoblotting to detect a specific Rho GTPase was performed as described above.

### Proteomics

Preparation of tissue lysates was performed as described above (in 200 uL of lysis buffer with 0.1M DTT) from pools of 8 sciatic nerves from P5 animals and using littermates (each biological replicate of CTR has a litter-matched biological replicate of RhoA cKO). Following proteolytic digestion with trypsin, peptides (corresponding to 500 ng protein/sample) were analyzed by LC-MS/MS and MS1-based label free quantitation was performed on a Hybrid Quadrupole-Orbitrap mass spectrometer (Q-Exactive, Thermo Fisher Scientific). Data was processed in Proteome Discoverer 2.4 using unique peptides only and applying the Minora algorithm for quantitative assessment to total peptide content. No imputation of missing values was performed and quantification channels were rejected if not found in more than 40% of biological replicates. A post-analysis refinement of the proteomics dataset was performed using the following criteria (Figure 3 A): an adjusted p value <0.050 (calculated by the ANOVA test and using the Benjamin Hochberg method, embedded in Proteome Discoverer) of the abundance ratio RhoA cKO/CTR value, at least two unique peptides identified/protein and PSM (peptide spectrum matches) value of at least 5. Gene ontology (GO) analysis was performed in g:Profiler (58). To predict cell type-predominant expression of proteins of interest, we searched the following scRNAseq databases: the sciatic nerve atlas (10) and neuron databases (22, 23). Detailed manual annotation of biological function/process was performed using Uniprot (59) and published literature.

### Antibodies, probes and dyes

For immunoblotting, primary and secondary antibodies were used as follows: mouse anti-GAPDH, HyTest 5G4, 1:20000; mouse anti-α-TUB, Sigma T5168, 1:5000; rabbit anti-RhoA, Cell Signaling 2117, diluted 1:1000; rabbit anti-RhoB, Cell Signaling 2098, diluted 1:500; rabbit anti-RhoC, Cell Signaling 3430, diluted 1:500; mouse anti-Rac1, Abcam ab33186, diluted 1:2000; rabbit anti-Cdc42, Cell Signaling 2466, diluted 1:1000; rabbit anti-OCT6 and rabbit anti-KROX20 (60), diluted 1:500, were a kind gift from D. Meijer, The University of Edinburgh, UK; rabbit anti-p-CFL1 (Ser3), Cell Signaling 3311, diluted 1:1000; mouse anti-CFL1, Abcam ab54532, diluted 1:1000; rabbit anti-p-PFN1 (Ser137), diluted 1:2000, was a kind gift from M. Diamond, UCSF, CA, USA; rabbit anti-PFN1, Abcam ab50667, diluted 1:1000; rabbit anti-p-MLC2 (Ser19), Cell Signaling 3671, diluted 1:500; rabbit anti-MLC2, Cell Signaling 8505, diluted 1:500; rabbit anti-p-ERK1/2 (Thr202/Tyr204), Cell Signaling 9101, diluted 1:2000; rabbit anti-ERK1/2, Cell Signaling 9102, diluted 1:2000; rabbit anti-p-S6 (Ser235/236), Cell Signaling 4858, diluted 1:1000, rabbit anti-S6, Cell Signaling 2317, diluted 1:1000; rabbit anti-PTEN, Cell Signaling 9188, diluted 1:1000; mouse anti-p-AKT (Ser473), Cell Signaling 4051, diluted 1:1000; rabbit anti-p-AKT (Thr308), Cell Signaling 2965, diluted 1:1000; rabbit anti-AKT, Cell Signaling 9272, diluted 1:1000; HRP goat anti-mouse, 115-035-146 and HRP donkey anti-rabbit, 711-035-152, from Jackson Immunoresearch. For immunoblotting, primary and secondary antibodies were used as follows: mouse anti-μ-TUB, Sigma T5168, 1:2000; goat anti-SOX10, RD Systems AF2864, diluted 1:250; mouse anti-ß3-Tubulin, SYSY 302302, diluted 1:2000; mouse anti-CASPR, NeuroMab 75-001, diluted 1:500; AF488 donkey anti-goat, ThermoFisher A-11055, diluted 1:1000; AF647 donkey anti-mouse, ThermoFisher A-31571, diluted 1:1000; AF488 goat anti-mouse, ThermoFisher A-11001, diluted 1:1000. Probes: AF594 Phalloidin, ThermoFisher A-12381, diluted 1:80; DAPI, ThermoFisher D1306, diluted 1:40000.

### Statistical analysis

Statistical analysis was performed using GraphPad Prism 7. Data distribution was tested for normality using the Shapiro-Wilk test. Comparisons between groups of measurements that follow a normal distribution and pass the F test to compare variances were performed using unpaired two-tailed Student’s t-test. For non-normal data, non-parametric tests were used. Statistical significance was assumed for p values <0.05.

## Author Contribution Statement

Study conception and design: A.I.S., J.B.R.; Data collection and analysis: A.I.S., M.R.G.M.; Interpretation of results: A.I.S., C.B., J.B.R.; Resources: C.B., J.B.R.; Manuscript preparation: A.I.S.; All authors reviewed the results and approved the final version of the manuscript.

## ACKNOWLEDGEMENTS

This work was funded by: FEDER - Fundo Europeu de Desenvolvimento Regional funds through the COMPETE 2020 - Operational Programme for Competitiveness and Internationalization (POCI), Portugal 2020, and by Portuguese funds through FCT - Fundação para a Ciência e Tecnologia/Ministério da Ciência, Tecnologia e Ensino Superior in the framework of the project PTDC/MED-NEU/31318/2017 (POCI-01-0145-FEDER-031318) awarded to J.B.R.; and Norte-01-0145-FEDER-000008 - Porto Neurosciences and Neurologic Disease Research Initiative, awarded to i3S and financed by NORTE 2020 through FEDER.

We thank Joana Paes de Faria and Jorge Pereira for their input during the project and final review of the manuscript. We thank the i3S core facilities for their support: Animal Facility, Advanced Light Microscopy and Histology and Electron Microscopy (members of PPBI, supported by POCI-01-0145-FEDER-022122), Proteomics (member of RNEM, supported by ROTEIRO/0028/2013; LISBOA-01-0145-FEDER-022125), and Cell Culture and Genotyping.

This document was prepared in Overleaf using the Henriques Lab bioRxiv template (https://henriqueslab.github.io/resources/bioRxivTemplate/).

---

**Figure S1.**
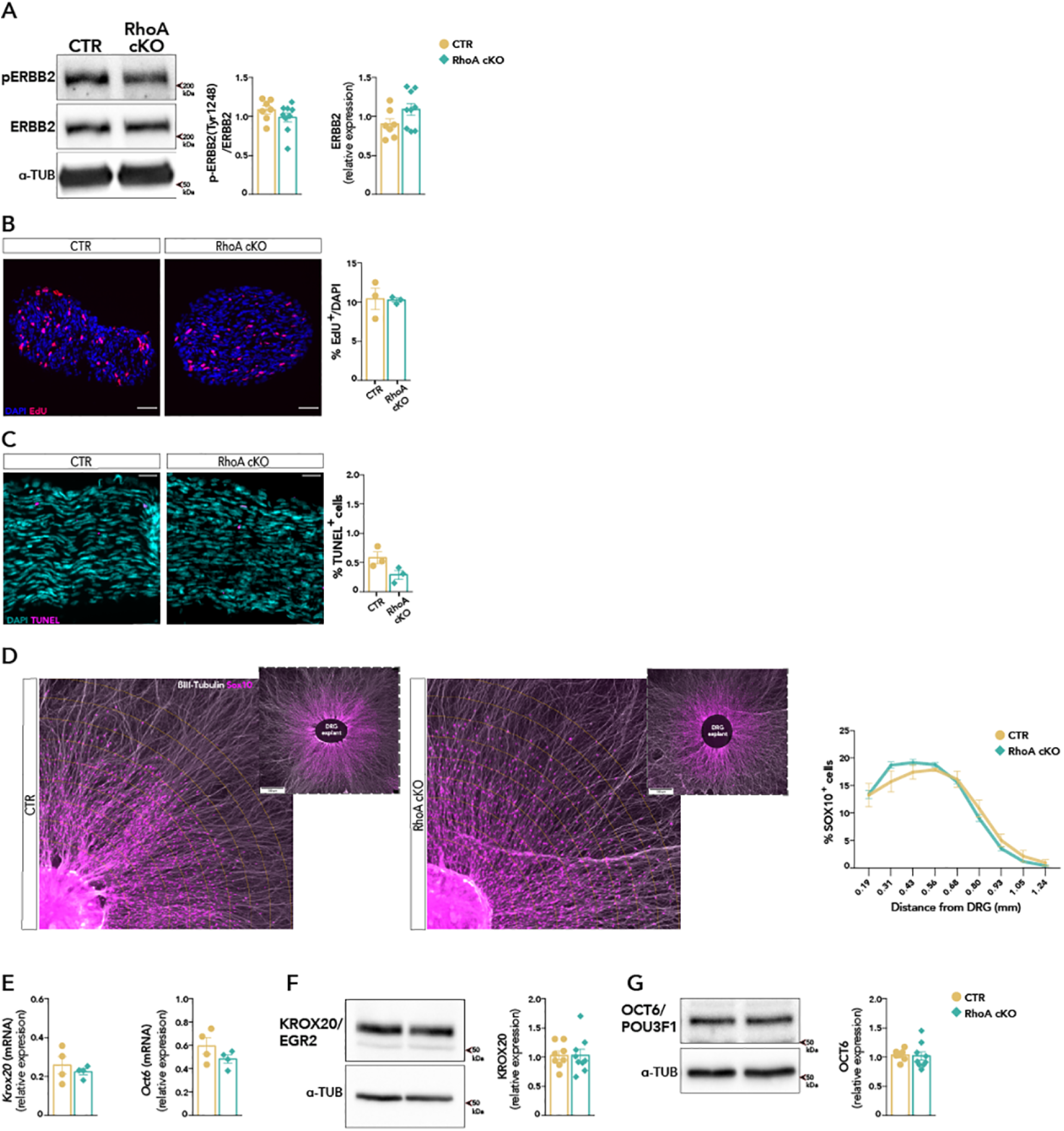
RhoA is dispensable for the proliferation, survival, and early differentiation of Schwann cells in postnatal sciatic nerves. **(A)** lmmunoblotting of phosphorylated and total ERBB2 in PS sciatic nerve lysates. Quantification of signal density relative to total protein or α-TUB.n=7 or 9 mice/group. **(B)** Representative images and quantification of IF analysis of cross sections of PS sciatic nerves. EdU (red) labels the nucleus (DAPI, blue) of proliferative cells. n=3 mice/ group. **(C)** Representative images and quantification of IF analysis of longitudinal sections of P5 sciatic nerves. TUNEL (magenta) labels apoptotic cells (nucleus in cyan, labeled with DAPI). n= 3 mice/group. **(D)** Analysis of Schwann cell migration at DIVS in DRG explants from E13.5 mouse embryos. SOX10 (magenta) labels Schwann cell nuclei and ßIII-Tubulin(grey) neurites. Ploted are the number of SCs as a function of distance to the DRG explant, quantified semi-automatically with a modified Sholl analysis(yellow lines). n=7/9 mice/ group. n.s. after multiple t-tests with Holm-Sidak correction. **(E-G)** Analysis of mRNA (E) and prot ein (F,G) levels of Krox20 and Oct6 by qRT-PCR and immunoblotting, respectively, of P5 sciatic nerve lysates. (E) Data represented as mean of relative expression (calculated as 2^−deltaC^) ± SEM. n= 4 mice/group. (F,G) Quantification of signal density relative to α-TUB. n= 8 or 9 mice/group. (A, B, C, D, E, F, G) Data presented as mean ± SEM and analyzed with unpaired t-test.

**Figure S2.**
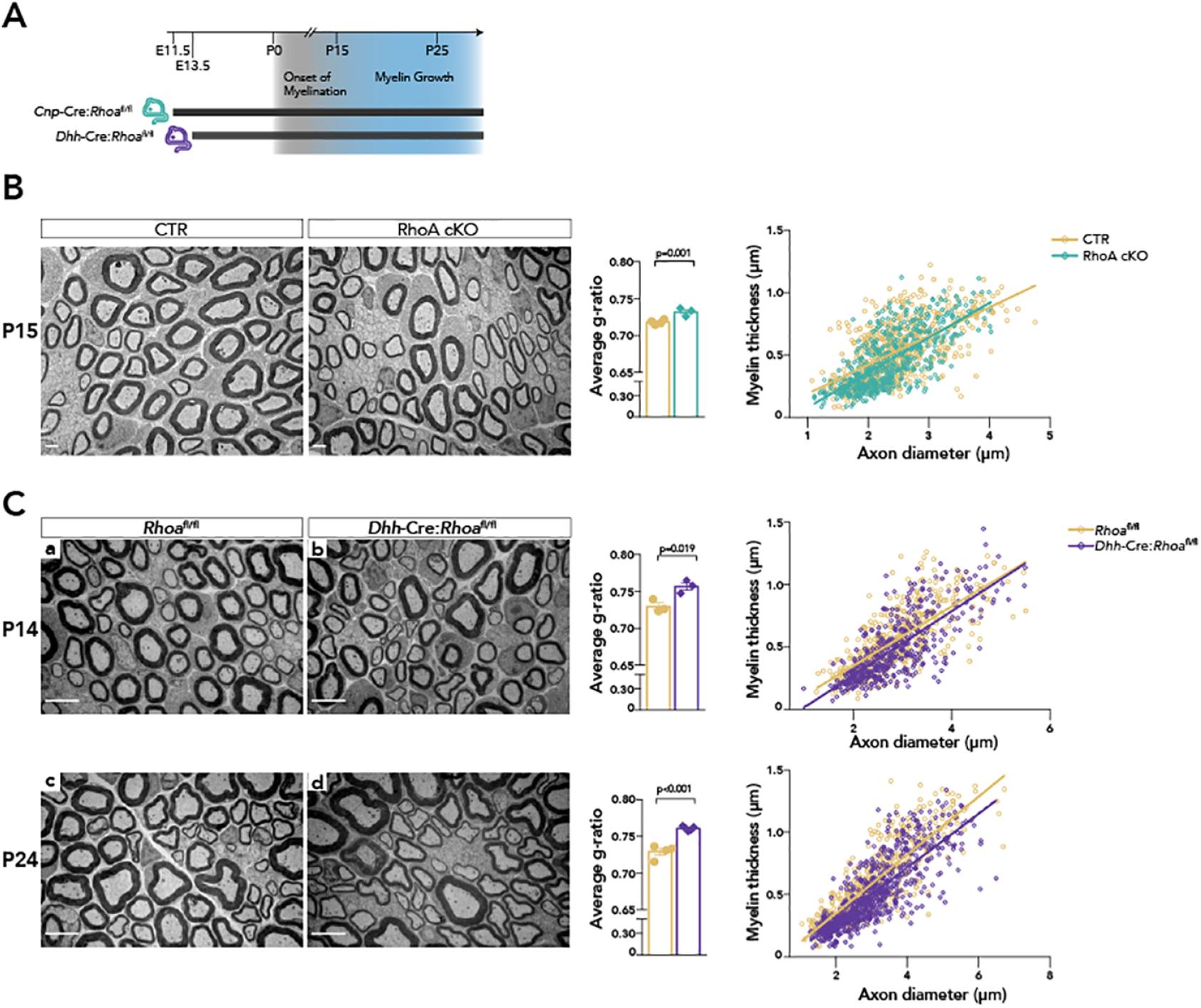
Myelinating cell-specific recombination of Rhoa delays development al myelination in the mouse PN S. **(A)** Schematic timeline of the genetic ablation of ***Rhoa*** specifically in myelinating cells or Schwann cells using the transgenic Cre recombinase-overexpressing strains *CnrrCre* and *Dhh-Cre,* respectively. **(B, C)** EM micrographs of sciatic nerve cross sections from CTR and RhoA cKO mice at P15 (B) and CTR and Dhh-Cre : *Rhoa^fl/fl^* (C) at P14 (a, b) and P24 (c,d). Bar graphs show average g-ratio and scatter plots show distribution of myelin sheath thickness as a function of axon caliber. At least 100 myelinated axons measured/animal. n=3 (B), n=3 or n=4/3 (C) mice/group. Data represented as mean ± SEM and analyzed with unpaired t-test. Scale bars are 2 μm.

**Figure S3.**
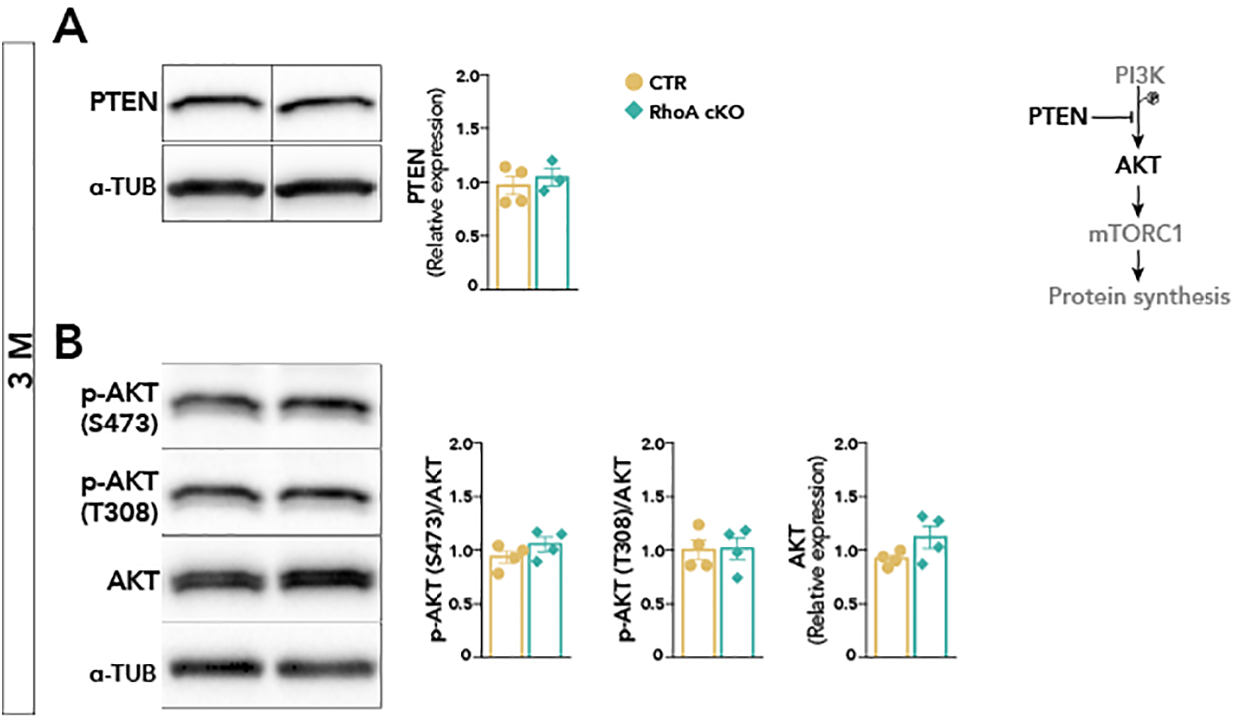
Activation of Akt signaling is unchanged in RhoA cKO hypermyelinated nerves. **(A)** lmmunoblotting of PTEN levels in sciatic nerves lysates at 3 months. Quantification of signal density relative to α-TUB. n=4 or 3 mice/group. **(B)** lmmunoblotting of phosphorylated and total AKT in sciatic nerve lysates at 3 months . Quantification of signal density relative to total protein or α-TUB. n=4 mice/group .

## Bibliography

1. M. L. Feltri, Y. Poitelon, and S. C. Previtali. How schwann cells sort axons: New concepts. Neuroscientist, 22(3):252–65, 2016. ISSN 1089-4098. doi: 10.1177/1073858415572361.

2. A. N. Muppirala, L. E. Limbach, E. F. Bradford, and S. C. Petersen. Schwann cell development: From neural crest to myelin sheath. Wiley Interdiscip Rev Dev Biol, 10(5):e398, 2021. ISSN 1759-7692. doi: 10.1002/wdev.398.

3. Yves Benninger, Tina Thurnherr, Jorge A. Pereira, Sven Krause, Xunwei Wu, Anna Chrostek-Grashoff, Dominik Herzog, Klaus-Armin Nave, Robin Franklin, Dies Meijer, Cord Brakebusch, Ueli Suter, and João B. Relvas. Essential and distinct roles for cdc42 and rac1 in the regulation of schwann cell biology during peripheral nervous system development. The Journal of Cell Biology, 177(6):1051–1061, 2007. ISSN 0021-9525. doi: 10.1083/jcb.200610108.

4. A. Nodari, D. Zambroni, A. Quattrini, F. A. Court, A. D’Urso, A. Recchia, V. L. Tybulewicz, L. Wrabetz, and M. L. Feltri. Beta1 integrin activates rac1 in schwann cells to generate radial lamellae during axonal sorting and myelination. J Cell Biol, 177(6):1063–75, 2007. ISSN 0021-9525. doi: 10.1083/jcb.200610014.

5. Jorge A. Pereira, Yves Benninger, Reto Baumann, Ana Gonçalves, Murat Özçelik, Tina Thurnherr, Nicolas Tricaud, Dies Meijer, Reinhard Fässler, Ueli Suter, and João B. Relvas. Integrin-linked kinase is required for radial sorting of axons and schwann cell remyelination in the peripheral nervous system. The Journal of Cell Biology, 185(1):147–161, 2009. ISSN 0021-9525. doi: 10.1083/jcb.200809008.

6. Laura Montani, Tina Buerki-Thurnherr, Joana P. de Faria, Jorge A. Pereira, Nuno G. Dias, Rui Fernandes, Ana F. Gonçalves, Attila Braun, Yves Benninger, Ralph T. Böttcher, Mercedes Costell, Klaus-Armin A. Nave, Robin J. Franklin, Dies Meijer, Ueli Suter, and João B. B. Relvas. Profilin 1 is required for peripheral nervous system myelination. Development, 141(7):1553–1561, 2014. ISSN 0950-1991. doi: 10.1242/dev.101840.

7. M. Amano, K. Chihara, K. Kimura, Y. Fukata, N. Nakamura, Y. Matsuura, and K. Kaibuchi. Formation of actin stress fibers and focal adhesions enhanced by rho-kinase. Science, 275 (5304):1308–11, 1997. ISSN 0036-8075. doi: 10.1126/science.275.5304.1308.

8. K. Kimura, M. Ito, M. Amano, K. Chihara, Y. Fukata, M. Nakafuku, B. Yamamori, J. Feng, T. Nakano, K. Okawa, A. Iwamatsu, and K. Kaibuchi. Regulation of myosin phosphatase by rho and rho-associated kinase (rho-kinase). Science, 273(5272):245–8, 1996. ISSN 0036-8075. doi: 10.1126/science.273.5272.245.

9. A. J. Ridley and A. Hall. The small gtp-binding protein rho regulates the assembly of focal adhesions and actin stress fibers in response to growth factors. Cell, 70(3):389–99, 1992. ISSN 0092-8674. doi: 10.1016/0092-8674(92)90163-7.

10. D. Gerber, J. A. Pereira, J. Gerber, G. Tan, S. Dimitrieva, E. Yanguez, and U. Suter. Transcriptional profiling of mouse peripheral nerves to the single-cell level to build a sciatic nerve atlas (snat). Elife, 10, 2021. ISSN 2050-084X. doi: 10.7554/eLife.58591.

11. S. D. Ackerman, R. Luo, Y. Poitelon, A. Mogha, B. L. Harty, M. D’Rozario, N. E. Sanchez, K. K. Lakkaraju, P. Gamble, J. Li, J. Qu, M. R. MacEwan, W. Z. Ray, A. Aguzzi, M. L. Feltri, X. Piao, and K. R. Monk. Gpr56/adgrg1 regulates development and maintenance of peripheral myelin. J Exp Med, 215(3):941–961, 2018. ISSN 1540-9538. doi: 10.1084/jem.20161714.

12. C. V. Melendez-Vasquez, S. Einheber, and J. L. Salzer. Rho kinase regulates schwann cell myelination and formation of associated axonal domains. J Neurosci, 24(16):3953–63, 2004. ISSN 1529-2401. doi: 10.1523/JNEUROSCI.4920-03.2004.

13. J. Wen, C. Qian, M. Pan, X. Wang, Y. Li, Y. Lu, Z. Zhou, Q. Yan, L. Li, Z. Liu, W. Wu, and J. Guo. Lentivirus-mediated rna interference targeting rhoa slacks the migration, proliferation, and myelin formation of schwann cells. Mol Neurobiol, 54(2):1229–1239, 2017. ISSN 1559-1182. doi: 10.1007/s12035-016-9733-5.

14. J. Wen, D. Tan, L. Li, X. Wang, M. Pan, and J. Guo. Rhoa regulates schwann cell differentiation through jnk pathway. Exp Neurol, 308:26–34, 2018. ISSN 1090-2430. doi: 10.1016/j.expneurol.2018.06.013.

15. C. Lappe-Siefke, S. Goebbels, M. Gravel, E. Nicksch, J. Lee, P. E. Braun, I. R. Griffiths, and K. A. Nave. Disruption of cnp1 uncouples oligodendroglial functions in axonal support and myelination. Nat Genet, 33(3):366–74, 2003. ISSN 1061-4036. doi: 10.1038/ng1095.

16. C. Taveggia. Schwann cells-axon interaction in myelination. Curr Opin Neurobiol, 39:24–9, 2016. ISSN 1873-6882. doi: 10.1016/j.conb.2016.03.006.

17. A. Piekny, M. Werner, and M. Glotzer. Cytokinesis: welcome to the rho zone. Trends Cell Biol, 15(12):651–8, 2005. ISSN 0962-8924. doi: 10.1016/j.tcb.2005.10.006.

18. L. Tavares, P. Gracio, R. Ramos, R. Traquete, J. B. Relvas, and P. S. Pereira. The pebble/rho1/anillin pathway controls polyploidization and axonal wrapping activity in the glial cells of the drosophila eye. Dev Biol, 473:90–96, 2021. ISSN 1095-564X. doi: 10.1016/j.ydbio.2021.02.002.

19. Y. Wang, H. L. Teng, and Z. H. Huang. Intrinsic migratory properties of cultured schwann cells based on single-cell migration assay. PLoS One, 7(12):e51824, 2012. ISSN 1932-6203. doi: 10.1371/journal.pone.0051824.

20. Fuzi Jin, Baoxia Dong, John Georgiou, Qiuhong Jiang, Jinyi Zhang, Arjun Bharioke, Frank Qiu, Silvia Lommel, Laura M. Feltri, Lawrence Wrabetz, John C. Roder, Joel Eyer, Xiequn Chen, Alan C. Peterson, and Katherine A. Siminovitch. N-wasp is required for schwann cell cytoskeletal dynamics, normal myelin gene expression and peripheral nerve myelination. Development, 138(7):1329–1337, 2011. ISSN 0950-1991. doi: 10.1242/dev.058677.

21. J. T. Velasquez, J. A. St John, L. Nazareth, and J. A. K. Ekberg. Schwann cell lamellipodia regulate cell-cell interactions and phagocytosis. Mol Cell Neurosci, 88:189–200, 2018. ISSN 1095-9327. doi: 10.1016/j.mcn.2018.01.001.

22. D. Usoskin, A. Furlan, S. Islam, H. Abdo, P. Lonnerberg, D. Lou, J. Hjerling-Leffler, J. Haeggstrom, O. Kharchenko, P. V. Kharchenko, S. Linnarsson, and P. Ernfors. Unbiased classification of sensory neuron types by large-scale single-cell rna sequencing. Nat Neurosci, 18 (1):145–53, 2015. ISSN 1546-1726. doi: 10.1038/nn.3881.

23. A. Zeisel, H. Hochgerner, P. Lonnerberg, A. Johnsson, F. Memic, J. van der Zwan, M. Haring, E. Braun, L. E. Borm, G. La Manno, S. Codeluppi, A. Furlan, K. Lee, N. Skene, K. D. Harris, J. Hjerling-Leffler, E. Arenas, P. Ernfors, U. Marklund, and S. Linnarsson. Molecular architecture of the mouse nervous system. Cell, 174(4):999–1014 e22, 2018. ISSN 1097-4172. doi: 10.1016/j.cell.2018.06.021.

24. J. Patzig, O. Jahn, S. Tenzer, S. P. Wichert, P. de Monasterio-Schrader, S. Rosfa, J. Kuharev, K. Yan, I. Bormuth, J. Bremer, A. Aguzzi, F. Orfaniotou, D. Hesse, M. H. Schwab, W. Mobius, K. A. Nave, and H. B. Werner. Quantitative and integrative proteome analysis of peripheral nerve myelin identifies novel myelin proteins and candidate neuropathy loci. J Neurosci, 31 (45):16369–86, 2011. ISSN 1529-2401. doi: 10.1523/JNEUROSCI.4016-11.2011.

25. S. B. Siems, O. Jahn, M. A. Eichel, N. Kannaiyan, L. M. N. Wu, D. L. Sherman, K. Kusch, D. Hesse, R. B. Jung, R. Fledrich, M. W. Sereda, M. J. Rossner, P. J. Brophy, and H. B. Werner. Proteome profile of peripheral myelin in healthy mice and in a neuropathy model. Elife, 9, 2020. ISSN 2050-084X. doi: 10.7554/eLife.51406.

26. S. J. Lord, K. B. Velle, R. D. Mullins, and L.K. Fritz-Laylin. Superplots: Communicating reproducibility and variability in cell biology. J Cell Biol, 219(6), 2020. ISSN 1540-8140. doi: 10.1083/jcb.202001064.

27. F. Meng and F. Sachs. Visualizing dynamic cytoplasmic forces with a compliance-matched fret sensor. J Cell Sci, 124(Pt 2):261–9, 2011. ISSN 1477-9137. doi: 10.1242/jcs.071928.

28. M. Amano, M. Ito, K. Kimura, Y. Fukata, K. Chihara, T. Nakano, Y. Matsuura, and K. Kaibuchi. Phosphorylation and activation of myosin by rho-associated kinase (rho-kinase). J Biol Chem, 271(34):20246–9, 1996. ISSN 0021-9258. doi: 10.1074/jbc.271.34.20246.

29. Nicklaus Sparrow, Maria E. Manetti, Marga Bott, Tiffany Fabianac, Alejandra Petrilli, Margaret L. Bates, Mary B. Bunge, Stephen Lambert, and Cristina Fernandez-Valle. The actin-severing protein cofilin is downstream of neuregulin signaling and is essential for schwann cell myelination. Journal of Neuroscience, 32(15):5284–5297, 2012. ISSN 0270-6474. doi: 10.1523/JNEUROSCI.6207-11.2012.

30. Haibo Wang, Ambika Tewari, Steven Einheber, James L. Salzer, and Carmen V. Melendez-Vasquez. Myosin ii has distinct functions in pns and cns myelin sheath formation. The Journal of Cell Biology, 182(6):1171–1184, 2008. ISSN 0021-9525. doi: 10.1083/jcb.200802091.

31. Z. Li, C. D. Aizenman, and H. T. Cline. Regulation of rho gtpases by crosstalk and neuronal activity in vivo. Neuron, 33(5):741–50, 2002. ISSN 0896-6273. doi: 10.1016/s0896-6273(02)00621-9.

32. H. Bagci, N. Sriskandarajah, A. Robert, J. Boulais, I. E. Elkholi, V. Tran, Z. Y. Lin, M. P. Thibault, N. Dube, D. Faubert, D. R. Hipfner, A. C. Gingras, and J. F. Cote. Mapping the proximity interaction network of the rho-family gtpases reveals signalling pathways and regulatory mechanisms. Nat Cell Biol, 22(1):120–134, 2020. ISSN 1476-4679. doi: 10.1038/s41556-019-0438-7.

33. M. Jaegle, M. Ghazvini, W. Mandemakers, M. Piirsoo, S. Driegen, F. Levavasseur, S. Raghoenath, F. Grosveld, and D. Meijer. The pou proteins brn-2 and oct-6 share important functions in schwann cell development. Genes Dev, 17(11):1380–91, 2003. ISSN 0890-9369. doi: 10.1101/gad.258203.

34. C. Berti, A. Nodari, L. Wrabetz, and M. L. Feltri. Role of integrins in peripheral nerves and hereditary neuropathies. Neuromolecular Med, 8(1-2):191–204, 2006. ISSN 1535-1084. doi: 10.1385/NMM:8:1-2:191.

35. E. Domenech-Estevez, H. Baloui, X. Meng, Y. Zhang, K. Deinhardt, J. L. Dupree, S. Einheber, R. Chrast, and J. L. Salzer. Akt regulates axon wrapping and myelin sheath thickness in the pns. J Neurosci, 36(16):4506–21, 2016. ISSN 1529-2401. doi: 10.1523/JNEUROSCI.3521-15.2016.

36. M. E. Sheean, E. McShane, C. Cheret, J. Walcher, T. Muller, A. Wulf-Goldenberg, S. Hoelper, A. N. Garratt, M. Kruger, K. Rajewsky, D. Meijer, W. Birchmeier, G. R. Lewin, M. Selbach, and C. Birchmeier. Activation of mapk overrides the termination of myelin growth and replaces nrg1/erbb3 signals during schwann cell development and myelination. Genes Dev, 28(3):290–303, 2014. ISSN 1549-5477. doi: 10.1101/gad.230045.113.

37. G. Figlia, D. Gerber, and U. Suter. Myelination and mtor. Glia, 66(4):693–707, 2018. ISSN 1098-1136. doi: 10.1002/glia.23273.

38. G. Figlia, C. Norrmen, J. A. Pereira, D. Gerber, and U. Suter. Dual function of the pi3k-akt-mtorc1 axis in myelination of the peripheral nervous system. Elife, 6, 2017. ISSN 2050-084X. doi: 10.7554/eLife.29241.

39. A. Ishii, M. Furusho, and R. Bansal. Sustained activation of erk1/2 mapk in oligodendrocytes and schwann cells enhances myelin growth and stimulates oligodendrocyte progenitor expansion. J Neurosci, 33(1):175–86, 2013. ISSN 1529-2401. doi: 10.1523/JNEUROSCI.4403-12.2013.

40. Dino P. Leone, Stéphane Genoud, Suzana Atanasoski, Reinhard Grausenburger, Philipp Berger, Daniel Metzger, Wendy B. Macklin, Pierre Chambon, and Ueli Suter. Tamoxifen-inducible glia-specific cre mice for somatic mutagenesis in oligodendrocytes and schwann cells. Molecular and cellular neurosciences. ISSN 1044-7431. doi: 10.1016/s1044-7431(03)00029-0.

41. P. Thomas, A. Pranatharthi, C. Ross, and S. Srivastava. Rhoc: a fascinating journey from a cytoskeletal organizer to a cancer stem cell therapeutic target. J Exp Clin Cancer Res, 38 (1):328, 2019. ISSN 1756-9966. doi: 10.1186/s13046-019-1327-4.

42. J. J. Bravo-Cordero, V. P. Sharma, M. Roh-Johnson, X. Chen, R. Eddy, J. Condeelis, and L. Hodgson. Spatial regulation of rhoc activity defines protrusion formation in migrating cells. J Cell Sci, 126(Pt 15):3356–69, 2013. ISSN 1477-9137. doi: 10.1242/jcs.123547.

43. E. M. Leitman, A. Tewari, M. Horn, M. Urbanski, E. Damanakis, S. Einheber, J. L. Salzer, P. de Lanerolle, and C. V. Melendez-Vasquez. Mlck regulates schwann cell cytoskeletal organization, differentiation and myelination. J Cell Sci, 124(Pt 22):3784–96, 2011. ISSN 1477-9137. doi: 10.1242/jcs.080200.

44. S. Niemann, M. W. Sereda, U. Suter, I. R. Griffiths, and K. A. Nave. Uncoupling of myelin assembly and schwann cell differentiation by transgenic overexpression of peripheral myelin protein 22. J Neurosci, 20(11):4120–8, 2000. ISSN 1529-2401.

45. N. Novak, V. Bar, H. Sabanay, S. Frechter, M. Jaegle, S. B. Snapper, D. Meijer, and E. Peles. N-wasp is required for membrane wrapping and myelination by schwann cells. J Cell Biol, 192(2):243–50, 2011. ISSN 1540-8140. doi: 10.1083/jcb.201010013.

46. Laurent Blanchoin, Rajaa Boujemaa-Paterski, Cécile Sykes, and Julie Plastino. Actin dynamics, architecture, and mechanics in cell motility. Physiological reviews, 94(1):235–263, 2014. ISSN 1522-1210. doi: 10.1152/physrev.00018.2013.

47. P. Chugh and E. K. Paluch. The actin cortex at a glance. J Cell Sci, 131(14), 2018. ISSN 1477-9137. doi: 10.1242/jcs.186254.

48. C. V. Melendez-Vasquez, J. C. Rios, G. Zanazzi, S. Lambert, A. Bretscher, and J. L. Salzer. Nodes of ranvier form in association with ezrin-radixin-moesin (erm)-positive schwann cell processes. Proc Natl Acad Sci U S A, 98(3):1235–40, 2001. ISSN 0027-8424. doi: 10.1073/pnas.98.3.1235.

49. P. W. Gunning, E. C. Hardeman, P. Lappalainen, and D. P. Mulvihill. Tropomyosin - master regulator of actin filament function in the cytoskeleton. J Cell Sci, 128(16):2965–74, 2015. ISSN 1477-9137. doi: 10.1242/jcs.172502.

50. M. A. El Azreq, M. Kadiri, M. Boisvert, N. Page, P. A. Tessier, and F. Aoudjit. Discoidin domain receptor 1 promotes th17 cell migration by activating the rhoa/rock/mapk/erk signaling pathway. Oncotarget, 7(29):44975–44990, 2016. ISSN 1949-2553. doi: 10.18632/oncotarget.10455.

51. M. W. Renshaw, D. Toksoz, and M. A. Schwartz. Involvement of the small gtpase rho in integrin-mediated activation of mitogen-activated protein kinase. J Biol Chem, 271(36): 21691–4, 1996. ISSN 0021-9258. doi: 10.1074/jbc.271.36.21691.

52. H. Hirata, M. Gupta, S. R. Vedula, C. T. Lim, B. Ladoux, and M. Sokabe. Actomyosin bundles serve as a tension sensor and a platform for erk activation. EMBO Rep, 16(2):250–7, 2015. ISSN 1469-3178. doi: 10.15252/embr.201439140.

53. B. Jackson, K. Peyrollier, E. Pedersen, A. Basse, R. Karlsson, Z. Wang, T. Lefever, A. M. Ochsenbein, G. Schmidt, K. Aktories, A. Stanley, F. Quondamatteo, M. Ladwein, K. Rottner, J. van Hengel, and C. Brakebusch. Rhoa is dispensable for skin development, but crucial for contraction and directed migration of keratinocytes. Mol Biol Cell, 22(5):593–605, 2011. ISSN 1939-4586. doi: 10.1091/mbc.E09-10-0859.

54. A. Ommer, G. Figlia, J. A. Pereira, A. L. Datwyler, J. Gerber, J. DeGeer, G. Lalli, and U. Suter. Ral gtpases in schwann cells promote radial axonal sorting in the peripheral nervous system. J Cell Biol, 218(7):2350–2369, 2019. ISSN 1540-8140. doi: 10.1083/jcb.201811150.

55. J. Schindelin, I. Arganda-Carreras, E. Frise, V. Kaynig, M. Longair, T. Pietzsch, S. Preibisch, C. Rueden, S. Saalfeld, B. Schmid, J. Y. Tinevez, D. J. White, V. Hartenstein, K. Eliceiri, P. Tomancak, and A. Cardona. Fiji: an open-source platform for biological-image analysis. Nat Methods, 9(7):676–82, 2012. ISSN 1548-7105. doi: 10.1038/nmeth.2019.

56. C. A. Schneider, W. S. Rasband, and K. W. Eliceiri. Nih image to imagej: 25 years of image analysis. Nat Methods, 9(7):671–5, 2012. ISSN 1548-7105.

57. K. R. Jessen and R. Mirsky. The origin and development of glial cells in peripheral nerves. Nat Rev Neurosci, 6(9):671–82, 2005. ISSN 1471-003X. doi: 10.1038/nrn1746.

58. U. Raudvere, L. Kolberg, I. Kuzmin, T. Arak, P. Adler, H. Peterson, and J. Vilo. g:profiler: a web server for functional enrichment analysis and conversions of gene lists (2019 update). Nucleic Acids Res, 47(W1):W191–W198, 2019. ISSN 1362-4962. doi: 10.1093/nar/gkz369.

59. Consortium UniProt. Uniprot: the universal protein knowledgebase in 2021. Nucleic Acids Res, 49(D1):D480–D489, 2021. ISSN 1362-4962. doi: 10.1093/nar/gkaa1100.

60. M. Ghazvini, W. Mandemakers, M. Jaegle, M. Piirsoo, S. Driegen, M. Koutsourakis, X. Smit, F. Grosveld, and D. Meijer. A cell type-specific allele of the pou gene oct-6 reveals schwann cell autonomous function in nerve development and regeneration. EMBO J, 21(17):4612–20, 2002. ISSN 0261-4189. doi: 10.1093/emboj/cdf475.

